# Discovering methylated DNA motifs in bacterial nanopore sequencing data with MIJAMP

**DOI:** 10.1101/2024.08.14.607972

**Authors:** Alyssa K. Tidwell, Evelyn Faust, Carrie A. Eckert, Adam M. Guss, William G. Alexander

**Affiliations:** Biosciences Division, Oak Ridge National Laboratory, Oak Ridge, TN 37830; The Bredesen Center for Interdisciplinary Research and Graduate Education, University of Tennessee-Knoxville, Knoxville, TN 37996; Department of Computational Medicine and Bioinformatics, University of Michigan, Ann Arbor, MI 48109

**Keywords:** Methylome, ONT sequencing, Restriction systems

## Abstract

Bacterial DNA methylation is involved in diverse cellular functions ranging from modulation of gene expression, DNA repair, and restriction-modification systems for defense against viruses and other foreign DNA. Methylome analysis determines sites within a bacterial chromosome that are methylated, revealing motifs that may be targeted by native restriction enzymes. Identification of these motifs is therefore critical to making an organism genetically tractable, where mimicking the methylome patterns in *Escherichia coli* allows plasmid DNA to be protected from restriction in the target organism and can therefore drastically enhance transformation efficiency. Oxford Nanopore Technologies (ONT) sequencing can detect methylated bases during sequencing, but software is needed to identify the corresponding methylated motifs within the data. Here, we develop MIJAMP (<u>M</u>IJAMP Is Just <u>A MethylBED P</u>arser), a software package that was developed to discover methylated motifs from the output of ONT’s Modkit or other data in the methylBED format. MIJAMP employs a human-driven refinement strategy that empirically validates all motifs against genome-wide methylation data, thus eliminating false, under-, or overexplained motifs. MIJAMP can also report methylation data on a specific, user-defined motif. Using MIJAMP, we determined the methylated motifs both in a control strain (wild type *E. coli*) and in *Picosynecococcus* sp. strain PCC7002, laying the foundation for improved transformation in this organism. MIJAMP is available at https://code.ornl.gov/alexander-public/mijamp/.

## Introduction

Like all other organisms, bacteria and archaea are subjected to cellular invasions by foreign genetic elements such as viruses and plasmids. Also like all other organisms, bacteria and archaea possess various adaptations to resist these invaders. One common mechanism is to encode restriction enzymes that hydrolyze foreign DNA at specific motifs. To prevent the restriction enzymes from degrading their own chromosomes, organisms methylate cytosine at the C5 (5-methylcytosoine, 5mC) or N4 atom (4-methylcytosine, 4mC) and/or adenine at the N6 atom (6-methyladenine, 6mA) in a system referred to as a restriction-modification, or R-M systems (1). Invading DNA will usually have different methylation patterns, and therefore will be targeted and destroyed by the endonuclease. R-M systems are frequently exchanged via horizontal gene transfer (2) and are typically hyper-variable between closely related strains. Therefore, understanding DNA methylation patterns in an organism requires methylome analysis for each targeted strain.

R-M systems are a barrier against genetic manipulation of their hosts, as the restriction nucleases will act on the heterologous DNA transformed into the cell because they do not share methylation patterns. Methods have been developed to evade these defenses to enable genetic manipulation, and many of these methods require knowledge of the motifs that are methylated. Approaches include removing restriction motifs from plasmid sequences (3) or re-creation of the host methylation patterns in a cloning strain of *Escherichia coli* (4,5). While a variety of these methylome analysis methods detect DNA methylation (6), two modern sequencing technologies are commonly used in this capacity: Pacific Biosciences (PacBio) and Oxford Nanopore Technologies (ONT). PacBio, long considered the gold-standard for long-read sequencing technologies, can detect 6mA and 4mC residues by a change in enzyme kinetics when the polymerase interacts with a methylated base. Unfortunately, 5mC modifications are only weakly detected by PacBio, thus requiring a prohibitive sequencing depth or chemical modification of the 5mC base for detection. In contrast, recent advances in ONT base-calling systems can detect all three mods equivalently. This advantage of ONT’s systems to detect all bacterial DNA modifications is highly valuable to investigators who require rapid and accurate methylation motif calls in novel bacterial species or strains.

While many software packages exist to extract and process DNA methylation from ONT sequencing runs (7,8,9), only a few are specialized to detect methylation motifs in bacterial genomes (10,11). These specialist packages have at least two common functions: some manner to call modified bases from the raw data, and a motif finder, usually software from the MEME suite (12). Variation exists in the strategies to achieve these functions. For instance, Nanodisco (10) compares the raw electrical signal of reads from methylated and unmethylated versions of the same genome to determine which bases are most likely modified. Nanodisco then extracts the genomic regions around these putative modified bases and inputs them into MEME, which finds patterns of enriched DNA motifs that surround these modified bases. MicrobeMod (11), in contrast, forgoes this raw current comparison method of modified base detection in favor of using the output from the ONT-provided DNA basecalling model. After the data are processed by Modkit, genomic regions around the modified bases are extracted and pattern detected using STREME. Unfortunately, many of these packages function as a type of black box system that takes an input and produces an output with no human intervention needed nor possible. This means investigators are taking at face value an automated analysis of potentially complex data with no ability to directly evaluate or alter the analysis. One package, Nanodisco, innovated a “motif refining” technique to empirically test each motif against the global dataset, which allows a user to correct mistakes made during motif detection and to clarify ambiguities arising from complex methylomes. However, Nanodisco is only compatible with data produced by older, out-of-production flowcells, and as of August 2024, no updates the Nanodisco code base have been made to correct this.

Thus, we aimed to create a simple, rapid, and interactive motif-finder that is compatible with ONT R10.4.1 flowcells. We developed MIJAMP (<u>M</u>IJAMP Is Just <u>A</u> M<u>ethylBED P</u>arser) to rapidly and accurately call methylation motifs in bacterial genomic DNA sequencing datasets. Like other packages, MIJAMP uses a single Modkit-parsed dataset per genome, but like Nanodisco it requires a user to manually refine and empirically validate every detected motif. We further discuss the MIJAMP workflow and demonstrate its capabilities on both simple and complex methylomes, as well as potential compatibilities with other sequencing technologies.

## Results

### The MIJAMP workflow

MIJAMP is a collection of executable scripts written in Python that act together to discover methylated motifs within prokaryotic genomes. These scripts rely on several Python packages along with Minimap (13), Modkit (github.com/nanoporetech/modkit), Dorado (https://github.com/nanoporetech/dorado), samtools (14), and MEME software present in the system’s PATH. The initial input files for MIJAMP are a BAM file containing modified base calls from Dorado and a reference genome in FASTA format. Next, the preprocess script maps the BAM file to the reference genome, processes and sorts the mapped result, and uses the final mapped, sorted BAM file as an input for Modkit. The output of Modkit (named “data.bed”) is saved into a working directory named by the user which includes the mapped, sorted BAM file, the reference genome, and any indexing files. After preprocessing has completed, the motif command is run to discover all DNA sequence motifs within that genome. First, the Modkit data are split into three pieces according to modification type (6mA, 4mC, 5mC) and filtered to remove low-quality base modification calls. The top 250 sites are extracted from the reference genome along with the 15 bases up and downstream from each site: these 31-mers are written to file and used as an input for MEME. Once complete, the MEME xml output is read and parsed to produce a list of unrefined motifs potentially present within the genome. For each motif in this list, a function calculates the location of the modification within the motif, and the motif and its modified base index are sent to a refining loop.

The purpose of the refining loop is to allow the user to empirically evaluate each putative motif against the genome-wide methylation (GWM, or the proportion of the motif that is called methylated across the genome) dataset; in other words, MEME is only a pattern finder and should be seen as the beginning to methylome discovery rather than the conclusion. By systematically producing all single nucleotide changes in a motif (both substitutions and indels), the user can experimentally validate the length and composition of said motif and substitute a better sequence in place of the one produced by MEME. The *E. coli* dataset included in the GitLab repository provides a new user an example of refinement in practice: in the second refinement loop of the 6mA data subset, MEME returns two putative motifs: (6mA)CNNNNNNGTGC and GC(6mA)CNNNNNNGTT. The former is immediately suspicious, as its GWM value is only 31.5%. This value is suggestive of a single missing base on either side of the motif: if 100% of a motif is methylated, then any terminal N-1 variant of that motif will be methylated approximately 25% of the time, with variation depending on specifics like positive selection for the site or GC content. Indeed, in the extended/retracted motif variant section, adding another A to the 5’ end of (6mA)CNNNNNNGTGC, producing A(6mA)CNNNNNNGTGC, brings the GWM value to 99.33%. Thus, a user can explore the data via a simple text interface to ensure that the motifs detected are as strenuously tested as possible. This refining loop can be executed as many times as the user desires until they are satisfied with the motif they have refined, and the sequence is saved to be reported later. After all motifs from MEME have been refined and either accepted or rejected, the data corresponding to the methylated base within the saved motifs are removed from the master dataset and a new loop is then initiated. This “refine then remove” strategy is inspired by and analogous to the refinement method used by Nanodisco and serves to better detect less-common motifs. Each dataset corresponding to a modification type is evaluated by these loops until no viable motifs are left to be found. At the end of this refinement process, the saved motifs of all types are written to an output tab-separated value file (summarized in Figure 1).

**Figure 1.**
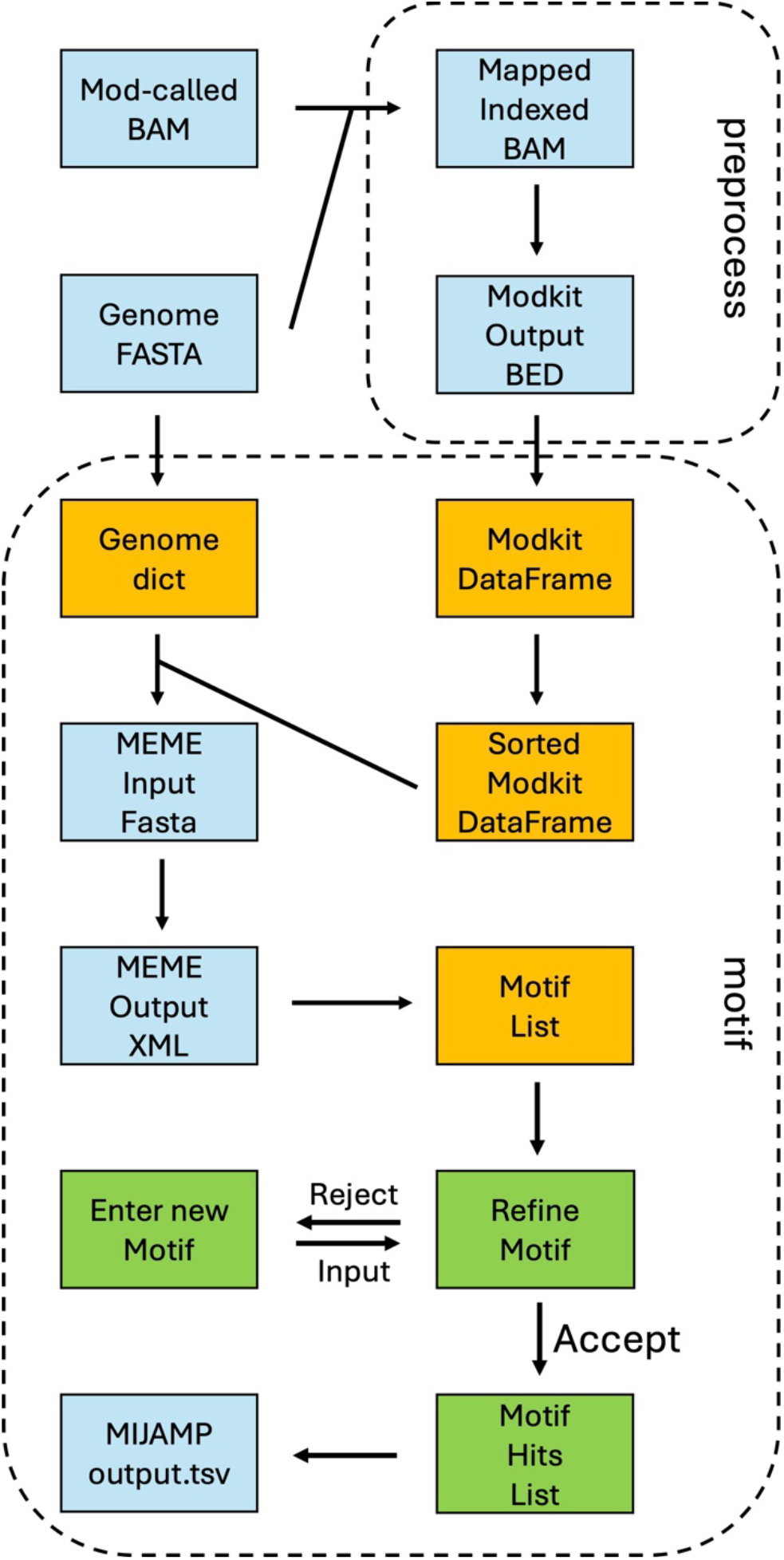
A flowchart summary of core MIJAMP functionality. Light blue boxes represent files present on the local system, gold boxes represent transient data structures, and green boxes represent the user refining loop feature of the software.

MIJAMP also includes two commands for exploring datasets: query and diagnostic. The query command accepts a user-defined motif and returns the number of methylated motifs, total number of motifs, and percent of motifs that are called as methylated, while the diagnostic command calculates the percentage of low-, mid-, and high-quality methylated base calls that are explained by the methylome derived from the motif command. In essence, query lets an investigator directly interrogate the methylome, while diagnostic provides a sanity check for an investigator nervous about potentially missing a motif.

### Benchmarking

MIJAMP and MicrobeMod were each run on the same datasets from *E. coli* K12 MG1655 and *Picosynecococcus sp*. PCC7002, which are included in the MIJAMP distribution. Each dataset was processed, and their outputs were compared using Dorado model v4.3.0 (Table S1) and v5.0.0 (Table 1). MIJAMP processed the *E. coli* and *Picosynecococcus* datasets in 315 and 956 seconds, respectively, while MicrobeMod completed its analysis in 1061 and 1410 seconds, respectively. The diagnostic script was then run using both methylomes in both organisms to determine how much methylation data they explain. To use the diagnostic script with the MicrobeMod methylomes, the motifs in each organism were placed in a spoofed output.tsv file one per line. As both MicrobeMod and MIJAMP predicted the same motifs in *E. coli*, which are also consistent with the known methylation patterns in this strain from the literature, they both explained the same proportion of methylated bases observed in the genome (Table 1). For *Picosynecococcus*, the MIJAMP methylome explained 99.9% of the 6mA data, 81.9% of the 5mC data, and 80.3% of the 4mC data, while the MicrobeMod methylome explained 94.3%, 65.8%, and 0% of the 6mA, 5mC, and 4mC data, respectively, as no 4mC motifs were returned.

**Table 1.**
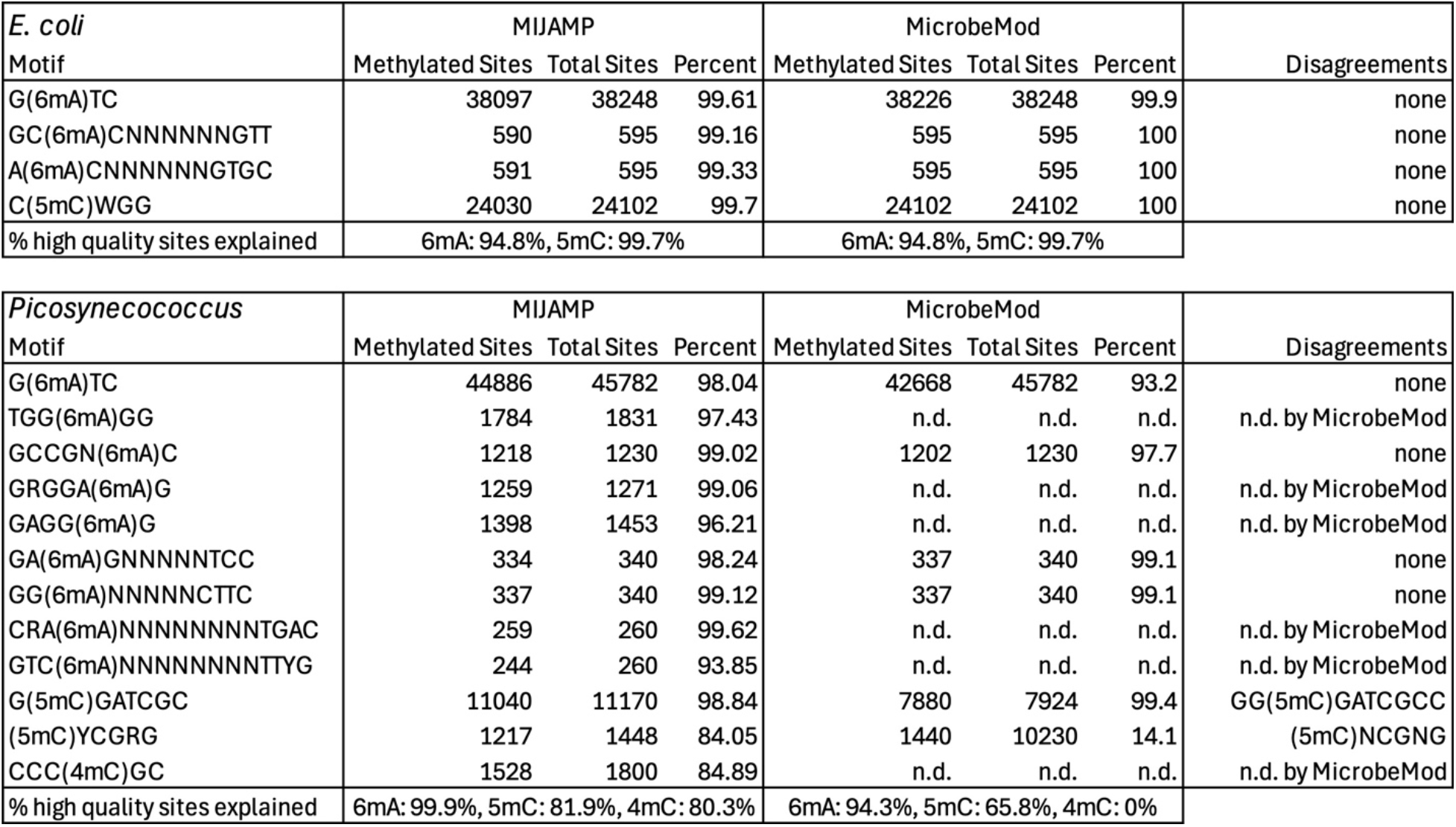
Benchmarking MIJAMP and MicrobeMod in conjunction with the ONT v5.0.0 modified base models. Modified bases are denoted in parentheses, n.d. = not detected.

While benchmarking used a dataset consisting of 470 Gbp worth of reads (or 100x) for each genome, we questioned what sequence depth minimum is required by MIJAMP. To explore these minimums, we made datasets containing 30x, 50x, 75x and 200x coverage worth of reads from the same master dataset as the *E. coli* dataset included in MIJAMP. Using the 30x dataset, MIJAMP was able to find correct motifs, though the GWM value for each was between 45 and 55%; this low result likely stems from a filtration step that removes low-coverage sites, which has a default value of 15 and, since the dataset contains data from both strands of DNA, we would expect a sizeable proportion of the data to be under this 15 read cutoff due to chance. This issue vanishes from every other read set. The only difference between the rest of the datasets is that the 50x dataset reported less-stringent motifs (i.e. RMACNNNNNNGYK instead of GCACNNNNNNGTT) than the deeper datasets. Thus, we recommend a target coverage of 100-200X, with 75X being an absolute minimum.

### MIJAMP and MicrobeMod are dependent on ONT methylation models

Initial development and benchmarking of MIJAMP used ONT’s v4.3.0 all-context modification models to identify which bases in the genome are modified. The methylomes resulting from using these models displayed unexpected methylation patterns in *Picosynecococcus*: while MIJAMP called a Dam methyltransferase site (G(6mA)TC) at a somewhat low GWM value (80%), MicrobeMod instead called that site as KG(6mA)TC. However, the real issue was the presence of the G(5mC)GATCGC motif alongside the G(6mA)TC motif, as the former completely enclosed the latter. By examining the Modkit output by hand, we observed that most G(5mC)GATCGC motifs lacked a 6mA residue at the 4^th^ base. This observation was confirmed via a bespoke script designed to compare methylation of two overlapping motifs, as only 22.9% of the GATC motifs inside a GCGATCGC motif were methylated, while 99.1% of the GATC motifs not inside this overlapping sequence were methylated. We reached out to ONT to inquire about model behavior when two different types of modifications were located close to one another, and their reply was that the v4.3.0 models may be reluctant to call one of the two modified bases in this scenario. They suggested using the newest v5.0.0 all-context models, which we used to regenerate the main dataset for benchmarking. Using the newer models appeared to have fixed the base calling interference issue from nearby modifications, but it did lead to some unexpected differences in the datasets. The newer models produced slightly lower GWM values in MIJAMP, though not substantially so. The motif composition improved with the newer models, where both software previously miscalled the (5mC)CYGRG motif as G(5mC)CCGGG (MIJAMP) or (5mC)CCGGG (MicrobeMod) with the older models. Using the newer models, MIJAMP called this motif correctly with 84% of sites called as methylated, while MicrobeMod called the motif (5mC)NCGNG with 14% or these sites called methylated.

## Discussion

Methylome discovery is a crucial early step in genetic tool development for the engineering of non-model bacteria. To that end, we developed MIJAMP to discover methylated motifs containing any of the three possible DNA modifications in bacteria from ONT sequencing. Additionally, the user-driven refinement method featured in MIJAMP helps to decrease the chance of miscalling the true sequence of a motif. MIJAMP provides utilities for both directly querying the dataset with a motif as well as confirming the extent of the data explained by a given methylome.

While MicrobeMod and MIJAMP called the *E. coli* methylome identically, the MIJAMP methylome explained substantially more data in *Picosynecococcus* sp. PCC7002 than MicrobeMod. This is likely due to the more complex methylome in *Picosynecococcus* sp. PCC7002, which contains seven 6mA, two 5mC, and one 4mC modified motifs. MIJAMP demonstrates its utility on these complex methylomes, as removing explained data from consideration by MEME helps prevent spurious motif calls, which in turn provides better starting motifs for refinement. Additionally, MIJAMP methylome calling explains a much higher proportion of the methylome data than MicrobeMod, as MicrobeMod did not call multiple well-supported motifs. Based on our results, we recommend using MIJAMP with a dataset representing at least 75x coverage of the target genome, though datasets larger than 100x are unlikely to improve the quality of the methylome produced.

MIJAMP is a methylBED file parser, and thus can hypothetically be used on any dataset written in this format. As the methylBED format is both well-defined and human readable, data obtained from other sequencing technologies, like Illumina bisulfite-Seq and PacBio SMRT-Seq, could likely be converted to the methylBED format for parsing by MIJAMP. We plan that future versions will provide options to directly query a methylBED file (and associated genome) for parsing.

## Methods

*E. coli* strain MG1655 was cultured in LB Miller medium overnight at 37 °C for genomic DNA extraction, which was done using the Zymo Research Quick-DNA Miniprep Plus Kit (Zymo Research, Irvine, CA). *Picosynecococcus* sp. strain PCC7002 was cultured in 20 mL glass bubble tubes in A+ medium (15) under a 50 µmol photons m^−2^ s^−1^ full-spectrum LED light for overnight culturing, then genomic DNA was extracted using the Zymo Research Quick-DNA Fungal/Bacterial Miniprep Kit. Each gDNA was used with the Ligation Sequencing Kit SQK-LSK114.24 (ONT, Oxford, UK) as instructed by ONT, except that the bead cleanup between the end repair and the barcode ligation steps was skipped. Sequencing was conducted using a MinION Mk. IIB run by an HP G5 Z8 workstation. Live basecalling was performed in Minknow v24.06.10 via Dorado v0.7.1 run on two NVIDIA RTX A6000 GPUs. Reference genomes were assembled from the obtained reads using the Trycycler v0.5.4 meta-assembler with default settings and suggested assemblers and accessories (Filtlong v0.2.1, Flye v2.9.4, Raven v1.5.3 miniasm/minipolish v0.3, and Medaka v1.5 (16-19)). Modified basecalling of the 100x coverage datasets by Dorado was performed on a separate HP G5 Z8 workstation, this one equipped with a single NVIDIA RTX A5000 GPU.

## Acknowledgements

We would like to thank Andrew Hren at CU-Boulder for providing a sample of gDNA from *Picosynecococcus sp*. PCC7002.

## Funding

This work was authored by Oak Ridge National Laboratory, which is managed by UT-Battelle LLC, for the U.S. DOE under contract DE-AC05-00OR22725. Funding was provided in part by the U.S. DOE, Office of Energy Efficiency and Renewable Energy Bioenergy Technologies Office (BETO) to the Agile BioFoundry. Funding was also provided in part by the DOE Office of Science Office of Biological and Environmental Research award DE-SC0023085.

## Author Contributions

Conceptualization: WGA, AMG, CAE; Code Development: WGA, AKT, EF; Sequence Data Generation: WGA; Writing: all authors; Project Administration: WGA, AMG, CAE.

## Competing Interests

The authors declare no competing interests.

## Data Availability

The data underlying this article are available in the MIJAMP GitLab repository (https://code.ornl.gov/alexander-public/mijamp/testData).

## Notes

https://code.ornl.gov/alexander-public/mijamp/

### Competing Interest Statement

The authors have declared no competing interest.

### Summary of Updates

Names, affiliations, and the GitLab URL were updated. Some ORCID numbers and an Acknowledgement section were added.

