## Supplementary Material for "Discovering methylated DNA motifs in bacterial nanopore sequencing data with MIJAMP"

| *E. coli* | MIJAMP | | | MicrobeMod | | | |
|---|---|---|---|---|---|---|---|
| Motif | Methylated Sites | Total Sites | Percent | Methylated Sites | Total Sites | Percent | Disagreements |
| G(6mA)TC | 38097 | 38248 | 99.61 | 38226 | 38248 | 99.9 | none |
| GC(6mA)CNNNNNNGTT | 590 | 595 | 99.16 | 595 | 595 | 100 | none |
| A(6mA)CNNNNNNGTGC | 591 | 595 | 99.33 | 595 | 595 | 100 | none |
| C(5mC)WGG | 24030 | 24102 | 99.7 | 24102 | 24102 | 100 | none |
| Run time | 315 sec | | | 1061 sec | | | |
| % high quality sites explained | 6mA: 94.8%, 5mC: 99.7% | | | 6mA: 94.8%, 5mC: 99.7% | | | |

| *Picosynecococcus* | MIJAMP | | | MicrobeMod | | | |
|---|---|---|---|---|---|---|---|
| Motif | Methylated Sites | Total Sites | Percent | Methylated Sites | Total Sites | Percent | Disagreements |
| G(6mA)TC | 36881 | 45782 | 80.56 | 19387 | 19390 | 99.9 | KG(6mA)TC |
| TGG(6mA)GG | 1821 | 1831 | 99.45 | 1788 | 1831 | 97.7 | none |
| GCCGN(6mA)C | 1221 | 1230 | 99.27 | n.d. | n.d. | n.d. | missing |
| GRGGA(6mA)G | 1261 | 1271 | 99.21 | n.d. | n.d. | n.d. | missing |
| GAGG(6mA)G | 1439 | 1453 | 99.04 | 1423 | 1487 | 95.7 | none |
| GA(6mA)GNNNNNTCC | 337 | 340 | 99.12 | 337 | 340 | 99.1 | none |
| GG(6mA)NNNNNCTTC | 338 | 340 | 99.41 | 337 | 340 | 99.1 | none |
| CRA(6mA)NNNNNNNNTGAC | 259 | 260 | 99.62 | n.d. | n.d. | n.d. | missing |
| GTC(6mA)NNNNNNNNTTYG | 240 | 260 | 92.31 | n.d. | n.d. | n.d. | missing |
| CG(6mA)TCGR | n.d. | n.d. | n.d. | 372 | 374 | 99.5 | not detected by MIJAMP |
| C**<u>N</u>**TCCYC | n.d. | n.d. | n.d. | 1772 | 4230 | 41.9 | not detected by MIJAMP |
| G(5mC)GATCGC | 11063 | 11170 | 99.04 | 7902 | 7924 | 99.7 | GG(5mC)GATCGCC |
| G(5mC)CCGGG | 336 | 359 | 93.59 | 900 | 990 | 90.9 | (5mC)CCGGG |
| CCC(4mC)GC | 1630 | 1800 | 90.56 | 1257 | 1801 | 69.8 | displayed as GCGGGG |
| Run time | 956 sec | | | 1410 sec | | | |
| % high quality sites explained | 6mA: 94.4%, 5mC: 92.5%, 4mC: 33.7% | | | 6mA: 55.9%, 5mC: 69.0%, 4mC: n.d. | | | |

Supplementary Table 1. MIJAMP and MicrobeMod outputs when using the v4.3.0 modified base models. Modified bases are denoted in parentheses, n.d. = not detected. The bolded, underlined base in the motif CNTCCYC is the listed location of the modified base by MicrobeMod.
